# Subregional Biomarkers in FDG PET for Alzheimer’s Diagnosis and Staging: An Interpretable and Explainable model

**DOI:** 10.1101/2024.12.27.630495

**Authors:** Ramin Rasi, Albert Guvenis, the Alzheimer’s Disease Neuroimaging Initiative

## Abstract

**Objective:** To investigate the radiomics features of the hippocampus and the amygdala subregions in FDG-PET images that can best differentiate Mild Cognitive Impairment (MCI), Alzheimer’s Disease (AD), and healthy patients.

**Methods:** Baseline FDG-PET data from 555 participants in the ADNI dataset were analyzed, comprising 189 cognitively normal (CN) individuals, 201 with MCI, and 165 with AD. The hippocampus and amygdala were segmented based on the DKT-Atlas, with additional subdivisions guided by probabilistic atlases from Freesurfer. Then radiomic features (n=120) were extracted from 38 hippocampal subregions and 18 amygdala nuclei using PyRadiomics. Various feature selection techniques, including ANOVA, PCA, Chi-square, and LASSO, were applied alongside nine machine learning classifiers.

**Results:** The Multi-Layer Perceptron (MLP) model combined with LASSO demonstrated excellent classification performance: ROC AUC of 0.957 for CN vs. AD, ROC AUC of 0.867 for MCI vs. AD, and ROC AUC of 0.782 for CN vs. MCI. Key regions, including the accessory basal nucleus, presubiculum head, and CA4 head, were identified as critical biomarkers. Features including GLRLM (Long Run Emphasis) and Small Dependence Emphasis (GLDM) showed strong diagnostic potential, reflecting subtle metabolic and microstructural changes often preceding anatomical alterations.

**Conclusion:** Specific hippocampal and amygdala subregions and their four radiomic features were found to have a significant role in the early diagnosis of AD, its staging, and its severity assessment by capturing subtle shifts in metabolic patterns. Furthermore, these features offer potential insights into the disease’s underlying mechanisms and model interpretability.

## **1** Introduction

### 1.1 Motivation

As Alzheimer’s disease (AD) continues to impact millions worldwide, the urgent need to unravel its complex mechanisms becomes increasingly pressing for both researchers and clinicians alike. Alzheimer’s disease is the most common form of dementia, accounting for 60-70% of all dementia cases (55.2 million people worldwide), with nearly 10 million new cases diagnosed annually, equating to one new case every 3 seconds [1]. Progression in Alzheimer’s includes complex neurobiological changes, including deposition of amyloid-beta plaques and tau tangles, along with pervasive inflammation. Each such factor plays a critical role in progressive cognitive decline [2].

Despite decades of dedicated research, the mechanisms underlying its progression from mild cognitive impairment (MCI) to advanced dementia remain poorly understood [3]. While certain brain regions are frequently implicated, no single region has been definitively linked to the disease’s progression [4]. Addressing these gaps necessitates a deeper exploration of understudied brain regions, network dysfunction, and metabolic alterations to identify novel biomarkers and therapeutic targets.

### 1.2 Importance of FDG PET

The diagnosis of AD by routine methods generally depends on a combination of cognitive assessments, biomarker studies, and neuroimaging [5], [6]. Nevertheless, early-stage diagnosis, in particular, is still highly challenging in the case of Mild cognitive impairment. MCI includes evident cognitive decline that might or might not, in fact, progress to AD; thus, it is highly important to determine which cases are most likely to progress. Imaging techniques allow us to investigate the brain changes associated with AD [7], [8]. Consequently, Fluorine-18-deoxyglucose positron emission tomography (FDG-PET) imaging has been invaluable in monitoring changes in brain metabolism indicative of neurodegeneration and provides early markers in cases of MCI and AD.

### 1.3 Importance of Hippocampus and Amygdala

Neuroimaging research has focused especially on the hippocampus and amygdala, two interconnected structures that AD notably affects and are involved in memory and emotional processing [9], [10]. In general, the structure composed of these two entities is referred to as the hippo-amygdala complex, which shows early signs of atrophy and metabolic changes associated with AD and, hence, has become pivotal in understanding the onset of the disease [11], [12]. The hippocampus is among the most frequently vulnerable regions in AD, where volume reductions have been observed, usually occurring together with memory loss. The amygdala is engaged in emotional memory and likewise showed degeneration associated with emotional symptoms in AD [13]. Damage at those specific locations has been shown to predict which MCI will progress to AD, hence underlining their potential as early diagnostic markers and targets for intervention [14].

### 1.4 Previous work on the role of subregions in AD

Recent studies emphasized the diagnostic potential of hippocampal analysis in AD and MCI. For example, [15] employed volumetric analysis of hippocampal and amygdala subfields using automated segmentation on MRI data, achieving high diagnostic accuracy in distinguishing AD from normal controls (AUC = 0.97) and moderate accuracy for MCI versus normal controls (AUC = 0.79) and AD versus MCI (AUC = 0.81). Similarly, [16] utilized functional connectivity analysis of the hippocampus with support vector machine learning on fMRI data, demonstrating accuracies exceeding 80% for all classification tasks, highlighting the role of abnormal hippocampal connectivity in differentiating stages of cognitive impairment.

Further advancements include [17] a lightweight 3D convolutional neural network model, which achieved an impressive AUC of 0.978 for AD versus normal controls using hippocampal MRI data, showcasing the strength of deep learning approaches in neuroimaging. [18] also achieved notable success with segmentation-based volumetric analysis of the hippocampus, reporting a classification accuracy of 94.4% for AD versus normal controls in a dataset of 373 participants.

### 1.5 The potential role of radiomics

Recent advances in radiomics, a computational approach to extracting quantitative features from medical images, offer deeper insights into brain changes that may indicate the progression of MCI to AD [19], [20]. Radiomics can systematically analyze textural, morphological, and intensity-based features of brain structures, often revealing subtle changes undetectable to the human eye [21]. This method shows particular promise for PET and MRI scans, as it enables detailed characterization of brain regions like the hippo-amygdala complex, where intricate structural and metabolic changes associated with AD emerge. Radiomics offers a powerful lens to explore brain imaging data beyond traditional atrophy measures, enabling a comprehensive view of region-specific texture and connectivity that may signal AD progression [22], [23]. For instance, texture-based radiomic features can highlight heterogeneity within a region, potentially reflecting cellular or metabolic variations characteristic of AD [24]. Radiomics research has identified texture and connectivity patterns in the hippocampus indicative of AD, suggesting that these could act as valuable markers for predicting cognitive decline [25]. Such analyses, however, can also potentially identify specific subfields or nuclei within these regions more susceptible to AD changes, which may be critical intervention points.

### 1.6 Objective of the study

While the hippocampus is widely recognized as an early AD target, newer research emphasizes the significance of examining subfields within this region, such as CA1, CA3, and the subiculum, which have unique vulnerabilities [26]. Likewise, the amygdala, though historically less studied, is increasingly acknowledged for its early structural and metabolic shifts in AD [27]. Therefore, these areas need to be studied in greater detail to open new potential avenues for targeted treatments focused on specific affected regions. This research study examines radiomic features within the subregions of the hippo-amygdala complex to enhance the understanding of their role in distinguishing stages of Alzheimer’s disease using FDG PET images. Since metabolic changes seen in FDG PET [28], [29] supersede anatomic changes, we can assume that these patterns of metabolic shifts may help in the early identification of AD and its transitions. Although some studies concentrated on finding radiomic biomarkers for AD using only FDG PET, to our knowledge, there has been no study that analyzes subfields and nuclei of the hippo-amygdala complex. As such, this is the first study that targets the metabolic properties of these areas in predicting MCI and AD. By combining FDG-PET data with radiomics, we aimed to uncover distinct metabolic features within the subregions of the hippocampus and amygdala that can differentiate MCI, AD, and healthy patients with more sensitivity, while offering an interpretable prediction algorithm.

## 2 Method

This study employed neuroimaging data from the Alzheimer’s Disease Neuroimaging Initiative (ADNI). After preprocessing and co-registering the FDG-PET and MRI images we segmented images into specific regions of interest. PyRadiomics was used to extract features from FDG-PET images, and dimensionality reduction techniques were applied to improve model performance. A diverse set of machine learning algorithms in combination with different feature selection methods were implemented and evaluated using stratified K-fold cross-validation. The outcomes were then refined and analyzed to support clinical decision-making, diagnosis, or prognosis. This approach enables the identification of informative biomarkers and the development of accurate diagnostic tools for early-stage cognitive impairment (Figure 1).

**Fig. 1.**
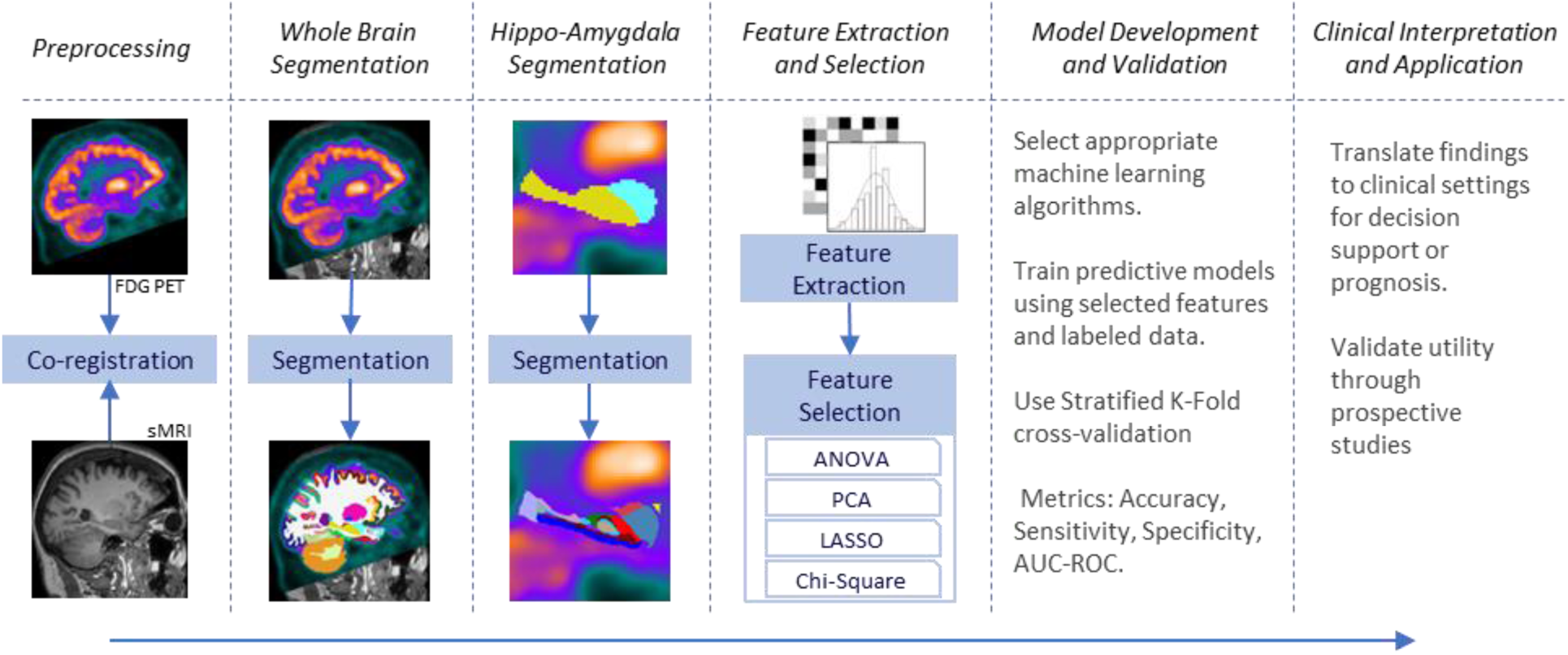
Pipeline for Feature Extraction and Classification from Hyppo-Amygdala Subregions.

### 2.1 ADNI and Participants

Data used in the preparation of this article were obtained from the Alzheimer’s Disease Neuroimaging Initiative (ADNI) database (adni.loni.usc.edu). The ADNI was launched in 2003 as a public-private partnership, led by Principal Investigator Michael W. Weiner, MD. The primary goal of ADNI has been to test whether serial magnetic resonance imaging (MRI), positron emission tomography (PET), other biological markers, and clinical and neuropsychological assessment can be combined to measure the progression of MCI and early Alzheimer’s disease. The data reveals a trend of decreasing cognitive function as measured by the Mini-Mental State Examination (MMSE) scores, with AD patients showing the lowest average score (23.1 ± 2.4) compared to MCI (27.5 ± 1.9) and CN (29.0 ± 1.2) groups, as shown in Table 1. This initial analysis suggests a potential link between clinical characteristics and disease progression, which our study aims to explore further using a radiomics approach.

**Table 1.** Clinical characteristics of participants.

| Clinical Diagnosis | No. Cases | Sex (M/F) | Age (mean ± SD) | MMSE (mean ± SD) |
| --- | --- | --- | --- | --- |
| AD | 165 | 93/72 | $74.6 \pm 8.1$ | $23.1 \pm 2.4$ |
| MCI | 201 | 121/80 | $72.7 \pm 7.4$ | $27.5 \pm 1.9$ |
| CN | 189 | 97/92 | $73.7 \pm 6.3$ | $29.0 \pm 1.2$ |
| Total | 555 | 306/249 | $73.6 \pm 7.3$ | $26.7 \pm 3.1$ |

### 2.2 Co-registration

To accurately co-register FDG-PET and sMRI images, we utilized the rigid transformation feature of the ANTsX Tool. This method relies exclusively on rotational and translational adjustments, ensuring that anatomical structures’ original shape and size remain unaltered. Rigid registration is particularly effective in scenarios where the images are only slightly misaligned in position or orientation, while their overall anatomical consistency is preserved. By employing this approach through ANTsX, we achieved precise and reliable co-registration of FDG-PET and sMRI data, facilitating seamless integration for subsequent analyses and interpretations [30].

### 2.3 FDG-PET Acquisition

For this study, we utilized FDG-PET data from the ADNI database. ADNI provides preprocessed FDG-PET images through four distinct pipelines; we selected the third pipeline, which includes realignment, co-registration to a standard template, spatial normalization, and smoothing. These comprehensive preprocessing steps, described in detail on the ADNI website (adni.loni.usc.edu), ensure uniformity across subjects, enabling robust comparative analyses. The dataset comprised dynamic 3D scans acquired 30–60 minutes after the administration of 185 MBq (5 mCi) of the radiotracer. Each scan consisted of six 5-minute frames. To facilitate precise comparisons of regional brain metabolic activity, the preprocessed data underwent Co-Reg, AVG, Standardized Image, and Voxel Size adjustments. This process standardized the images to a 160×160×96 voxel grid with 1.5 mm isotropic resolution, ensuring a consistent framework for subsequent analyses [31].

### 2.4 MRI Acquisition

MRI data was collected following standardized protocols designed to maintain consistency across subjects and studies. Imaging was conducted on high-field MRI scanners (3T or higher), utilizing a range of pulse sequences such as T1-weighted, T2-weighted, fluid-attenuated inversion recovery (FLAIR), and diffusion tensor imaging (DTI). To reduce variability between scanners, uniform pulse sequences and standardized imaging parameters—including slice thickness, field of view, echo time, and repetition time—were carefully implemented [32].

### 2.5 Segmentation

To segment the hippocampus and amygdala into their respective subfields and subnuclei, a two-step methodology was employed, blending broad anatomical mapping with fine-grained structural analysis.

The first step involved segmenting the entire brain into 95 distinct regions of interest (ROIs) using the Desikan-Killiany-Tourville (DKT) atlas within the FastSurfer pipeline. This automated process, powered by a deep learning-based approach, achieved segmentation with remarkable speed, significantly reducing processing time compared to traditional methods. FastSurfer provided a comprehensive anatomical overview, establishing a foundational spatial context for subsequent focused studies. By capturing major brain landmarks efficiently and accurately, this segmentation ensured a robust and time-efficient starting point for downstream analyses.[33].

In the second step, the hippocampus and amygdala were segmented using FreeSurfer. The hippocampus was delineated into 19 subfields for each hemisphere, encompassing the hippocampus proper (CA regions), subiculum, parasubiculum, and presubiculum. Similarly, the amygdala was divided into 9 subnuclei per hemisphere, reflecting its detailed internal architecture. FreeSurfer used probabilistic atlases and shape models to achieve segmentation and volume measurement [34], [35], [36]. The IDs for these subregions and subnuclei, referred to as FS IDs, are listed in Table 2.

**Table 2.**
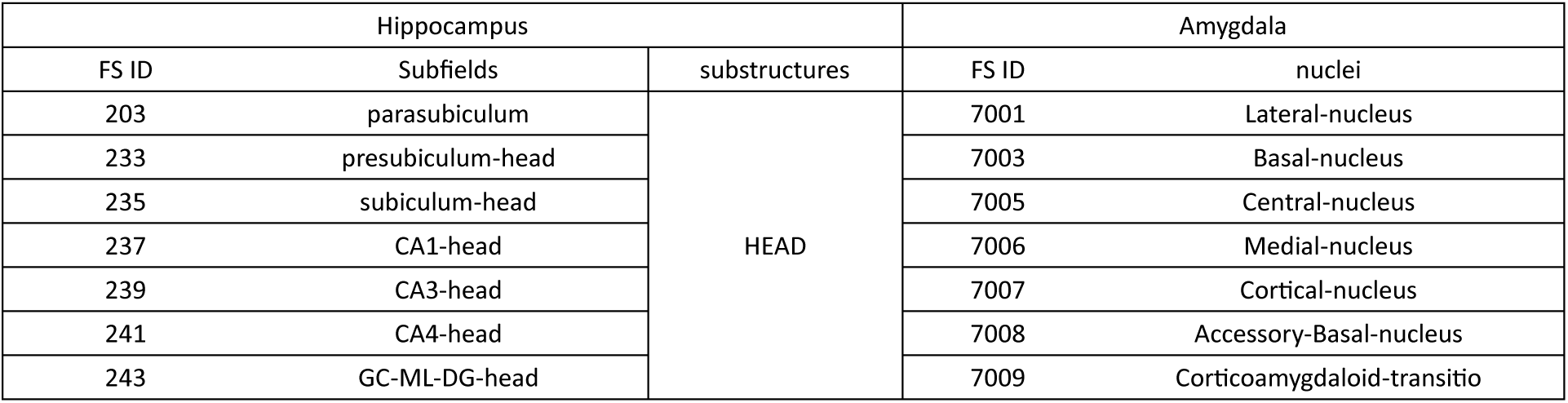
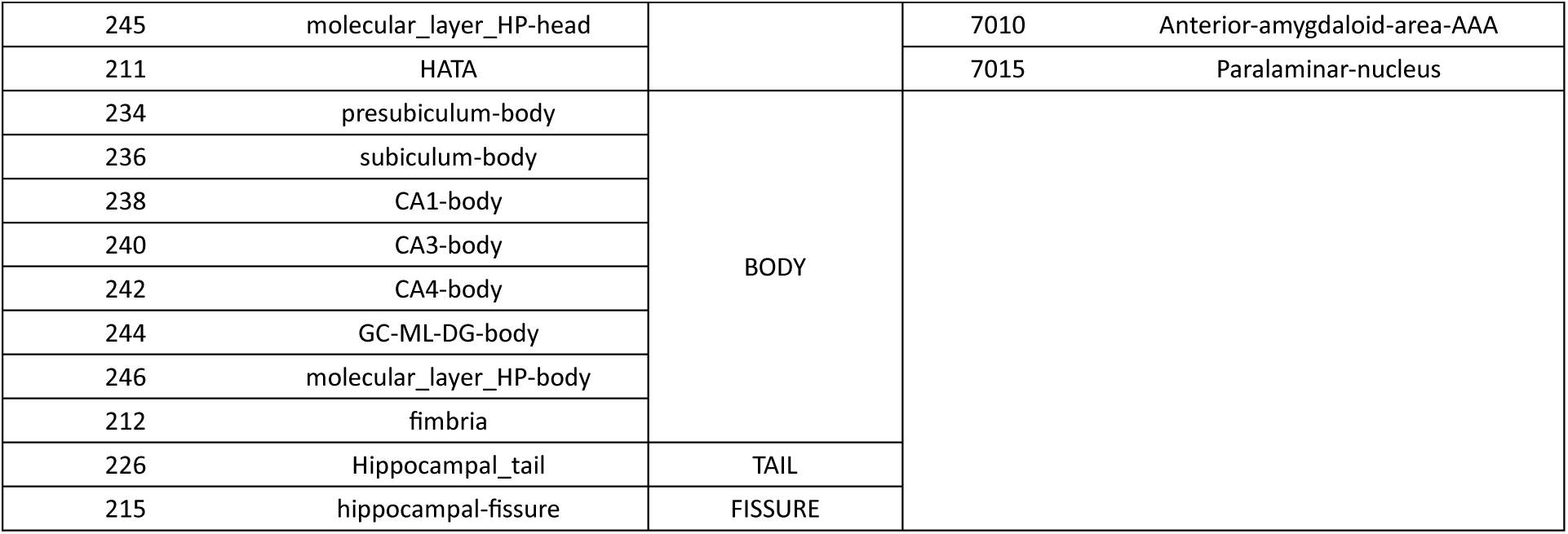
IDs and corresponding subregions of the hippocampus and nuclei of the amygdala as segmented by FreeSurfer, detailing the hierarchical organization and anatomical substructures.

### 2.6 Feature Extraction and Selection

To extract meaningful features from FDG-PET images, the PyRadiomics package was utilized to compute 120 feature classes across 56 predefined regions of interest (ROIs). These features spanned various metrics, including first-order statistics, texture features such as GLCM, GLRLM, NGTDM, and GLSZM, as well as shape-based characteristics [37].

To address the curse of dimensionality and enhance model performance, we examined different dimension reduction methods. Several techniques were explored, including filter-based approaches like ANOVA and Chi-square, along with embedded methods such as the Least Absolute Shrinkage and Selection Operator (LASSO). Furthermore, Principal Component Analysis (PCA) was employed to reduce the dimensionality of the feature space while retaining critical information. This systematic selection process aimed to identify the most predictive features, ultimately strengthening the performance of the resulting models.

For LASSO, a pipeline was constructed consisting of mean imputation, feature standardization, and the application of the LASSO model. The pipeline was optimized using grid search cross-validation over a range of regularization strengths alpha from 0.001 to 0.1 in 0.001 increments, with a 10-fold stratified cross-validation using the ROC AUC metric. This approach effectively identified the most influential features by shrinking less important coefficients to zero.

For PCA, the data was first standardized, and dimensionality reduction was performed by retaining the top 50 principal components, which captured the maximum variance in the dataset. A Random Forest classifier was then trained on the PCA-transformed data to assess feature importance within the reduced feature space.

In the ANOVA, the FF-statistic and corresponding p-values were calculated using the f_classif method, evaluating the linear relationship between each feature and the target variable. Features with higher FF values were considered more predictive.

In Chi-square, data was scaled using Min-Max normalization to satisfy the non-negative input requirement. The SelectKBest method was used to identify the top 50 features based on their Chi-square scores, which measure the dependency between features and the target variable.

### 2.7 Classification

We implemented a diverse array of machine learning classifiers to achieve a reliable evaluation. The classification framework incorporated algorithms spanning multiple methodological categories, including ensemble methods (Gradient Boosting (GB), Random Forest (RF), AdaBoost (AB)), decision-based techniques (Decision Tree (DT)), probabilistic models (Gaussian Naive Bayes (GNB)), kernel-based approaches (Gaussian Process (GP)), neural networks (Multi-layer Perceptron (MLP)), discriminant analysis (Quadratic Discriminant Analysis (QDA), and proximity-based methods (K-Nearest Neighbors (KNN)) [38]. To ensure robust evaluation, we employed stratified K-fold cross-validation. We computed key metrics such as ROC AUC, accuracy, sensitivity, and specificity for each fold to assess the model performance. We tuned the hyperparameters for each classifier using a grid search approach. This involved exploring a range of values for key parameters such as learning rate, number of estimators, maximum depth, and others, as shown in Table 3.

**Table 3.**
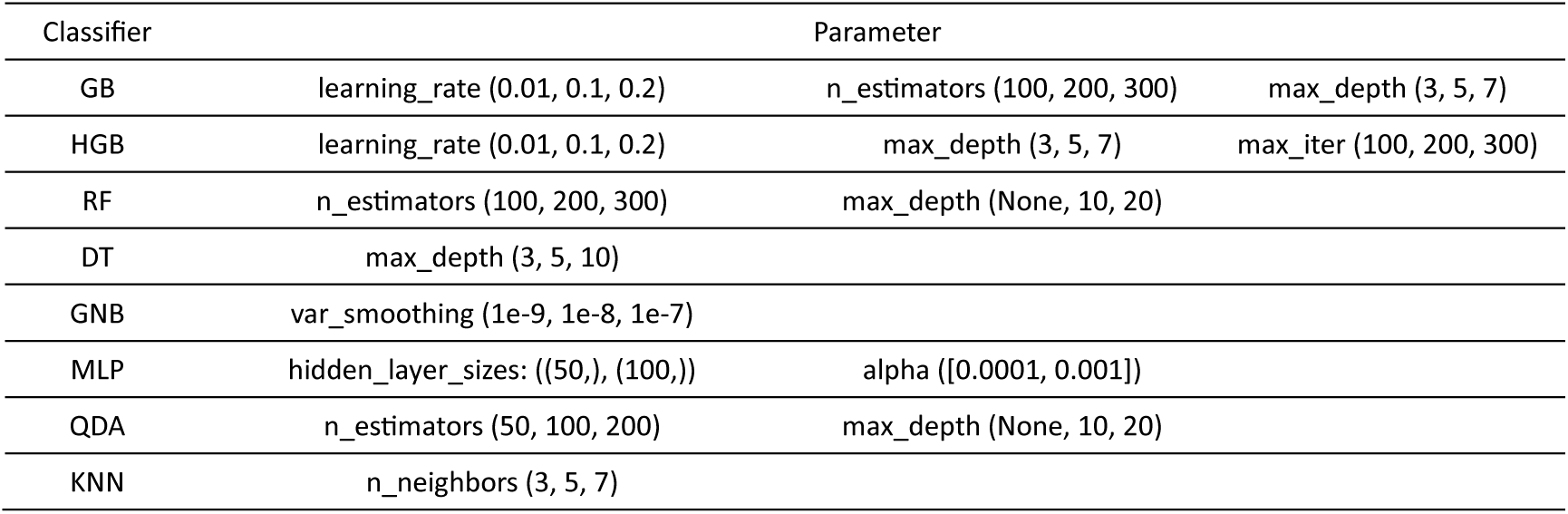
Values of the Hyperparameters for the Classifiers.

## 3 Result

### 3.1 Evaluation of Classifiers and Feature Selection Methods

Given the interdependence between feature selection methods and classifier performance, we evaluated multiple combinations of feature selection techniques including ANOVA, PCA, LASSO, Chi-square, and classifiers including GB, RF, AB, DT, GNB, GP, MLP, QDA, KNN to optimize diagnostic accuracy. Specifically, we applied these feature selection methods to radiomic features from two diagnostic groups: CN vs. AD and MCI vs. AD. Following feature reduction, each feature subset was tested across nine distinct classifiers. Classifier optimization was performed using grid search (cv=3) to fine-tune the hyperparameters as listed in Table 3. We employed Stratified K-Fold cross-validation (k=5) to split the dataset into balanced folds, ensuring each fold maintained the same proportion of AD, MCI, and CN participants as in the original dataset. Each classifier was trained and validated five times once on each fold, with one fold held out for testing and the remaining four used for training thus maximizing the stability and generalizability of model performance across the dataset.

We assessed model performance based on ROC AUC and accuracy, presented in Table 4, to determine the optimal feature selection and classifier combinations. For distinguishing CN from AD, combining the MLP classifier with the LASSO feature selection yielded the best ROC AUC scores of 0.957 and an accuracy of 0.901. In distinguishing MCI from AD, the MLP classifier paired with LASSO achieved the best AUC score of 0.867 and an accuracy of 0.77.

**Table 4.**
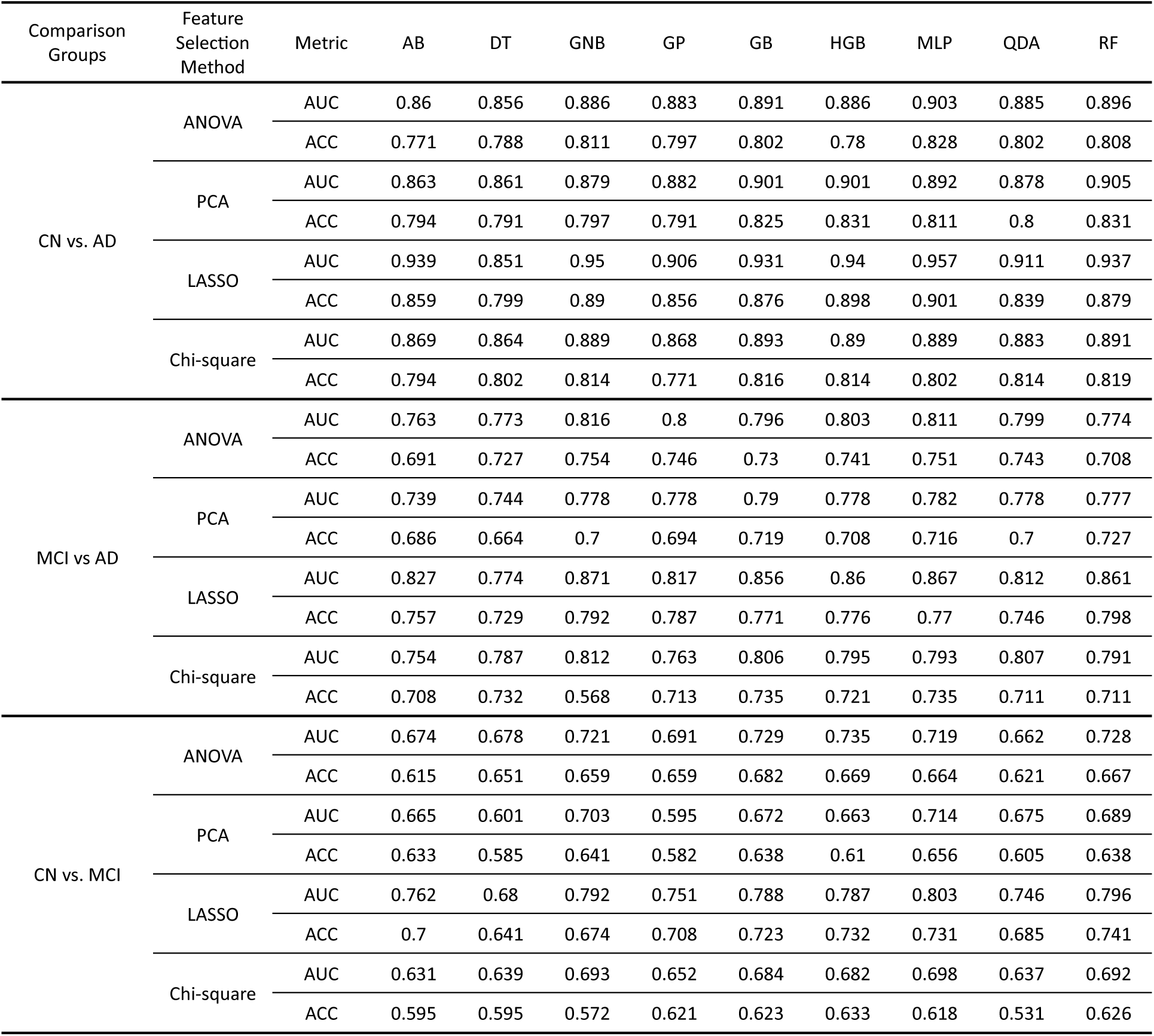
Classification Performance of Different Feature Selection Methods and Classifiers for AD and MCI vs CN.

We conducted an analysis of the top-performing models to investigate further the underlying radiomic features and subregions contributing to model performance. This analysis focused on the feature subsets and classifiers that were most effective for each comparison group (CN vs. AD and MCI vs. AD), enabling a detailed exploration of potential imaging biomarkers within hippocampal and amygdala subregions.

### 3.2 CN vs. AD

The comparative analysis of CN vs. AD classification was performed to identify the most predictive feature subset using the MLP classifier. A rank-based approach was applied, systematically evaluating all subsets of the top 50 features identified through the LASSO method. Features were added incrementally in each evaluation step, following their ranking based on co-efficiency values. For each subset, four metrics ROC AUC, accuracy, specificity, and sensitivity were computed. The aim was to determine the feature subset achieving the highest average of ROC AUC and accuracy.

As shown in Figure 2, the optimal subset comprised 36 features, where the MLP classifier demonstrated outstanding performance, achieving a ROC AUC of 0.951, an accuracy of 0.907, a specificity of 0.916, and a sensitivity of 0.897. This configuration was selected as the optimal point based on the average of ROC AUC and accuracy, emphasizing the significance of these metrics in evaluating prediction models. These results demonstrate the effectiveness of this subset in differentiating between AD and CN subjects.

**Fig. 2.**
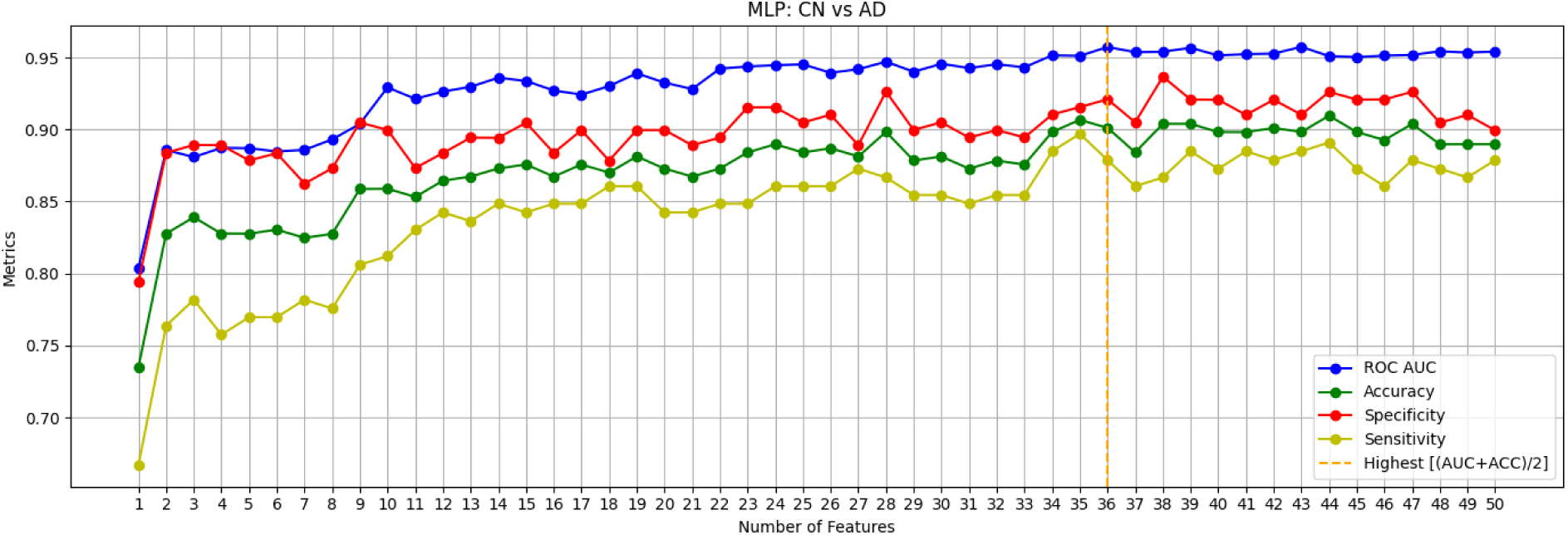
Performance of MLP Classifier with LASSO-Selected Features for CN vs. AD Discrimination.

To refine the model and achieve a more interpretable feature set, we applied Pearson correlation analysis to all the top 36 selected features (Figure 3). This step aimed to identify and reduce redundancy among the features by selecting those with the least interdependence. By focusing on uncorrelated features, we ensure that the selected variables provide unique contributions to the model, avoiding overlap in the information they represent.

**Fig. 3.**
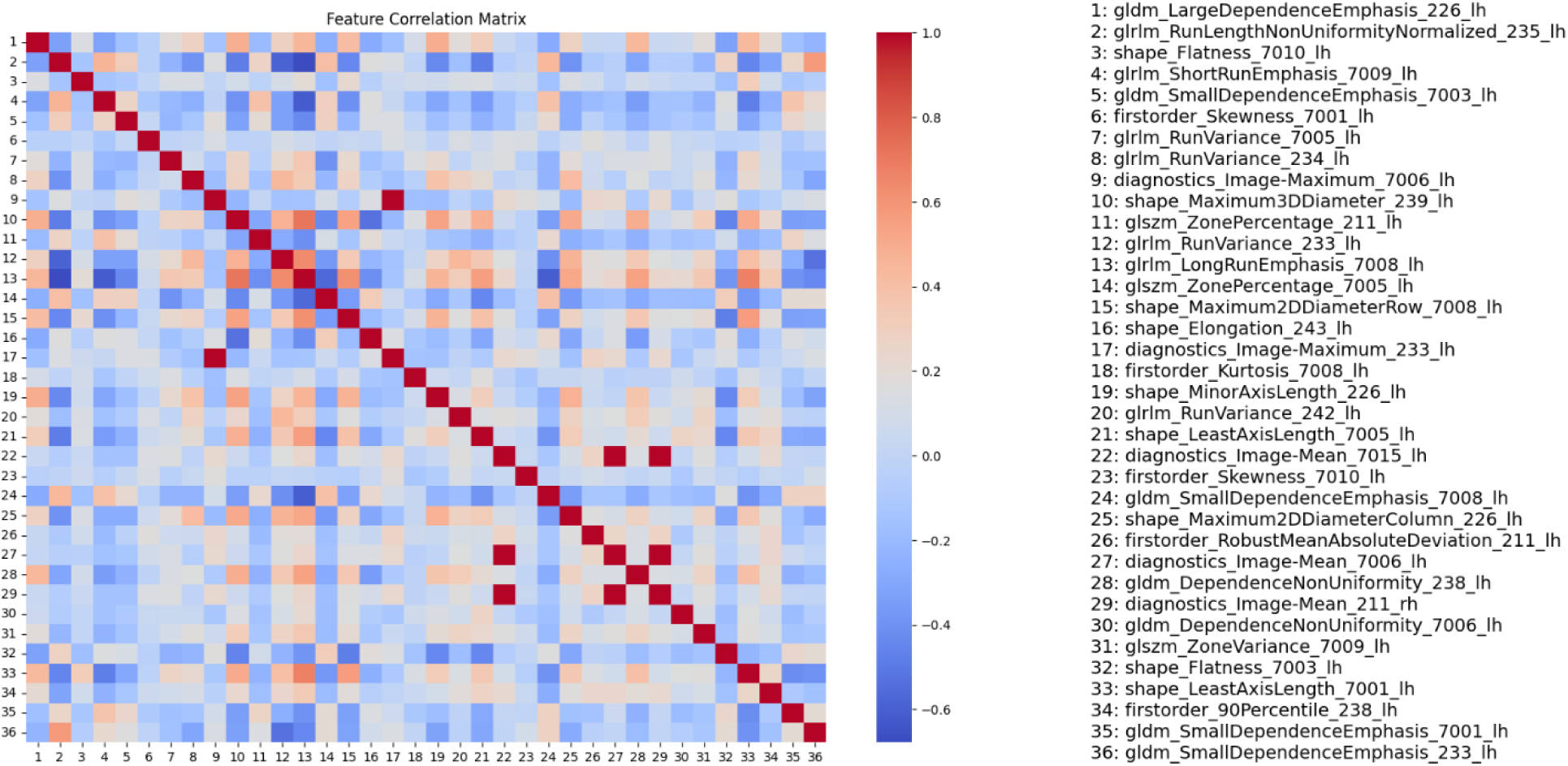
Feature correlation matrix for the group of features with the highest accuracy in discriminating CN from AD.

To identify a refined feature set with minimal redundancy and diverse representation, we selected the feature pairs with the lowest absolute correlation coefficients based on Pearson correlation analysis, indicating weak or negligible relationships. To validate these pairs, we further calculated p-values for their correlation coefficients using the Spearman rank correlation test, ensuring statistical confirmation of the correlations’ strength and significance while ruling out random variation. The resulting Table 5 summarizes the selected uncorrelated feature pairs, along with their correlation coefficients and corresponding p-values, providing a robust foundation for feature selection.

**Table 5:**
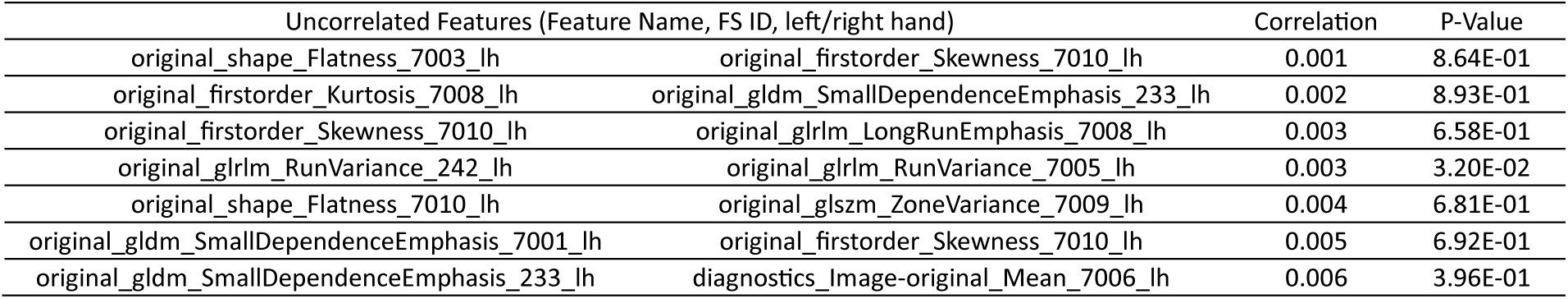
Highly uncorrelated feature pairs for distinguishing CN vs. AD identified through the Pearson correlation test, showing minimal correlation and corresponding p-values.

### 3.3 MCI vs AD

To identify the most predictive feature subset for differentiating Alzheimer’s disease AD from MCI, we assessed the performance of the MLP classifier using four metrics: ROC AUC, Accuracy, Specificity, and Sensitivity. These metrics were calculated for subsets of the top 50 LASSO-selected features ranked by a systematic method. The analysis revealed that the optimal feature subset consisted of 33 features, achieving an MLP classification performance of ROC AUC: 0.861, Accuracy: 0.806, Specificity: 0.821, Sensitivity: 0.788, as presented in Figure 4.

**Fig. 4.**
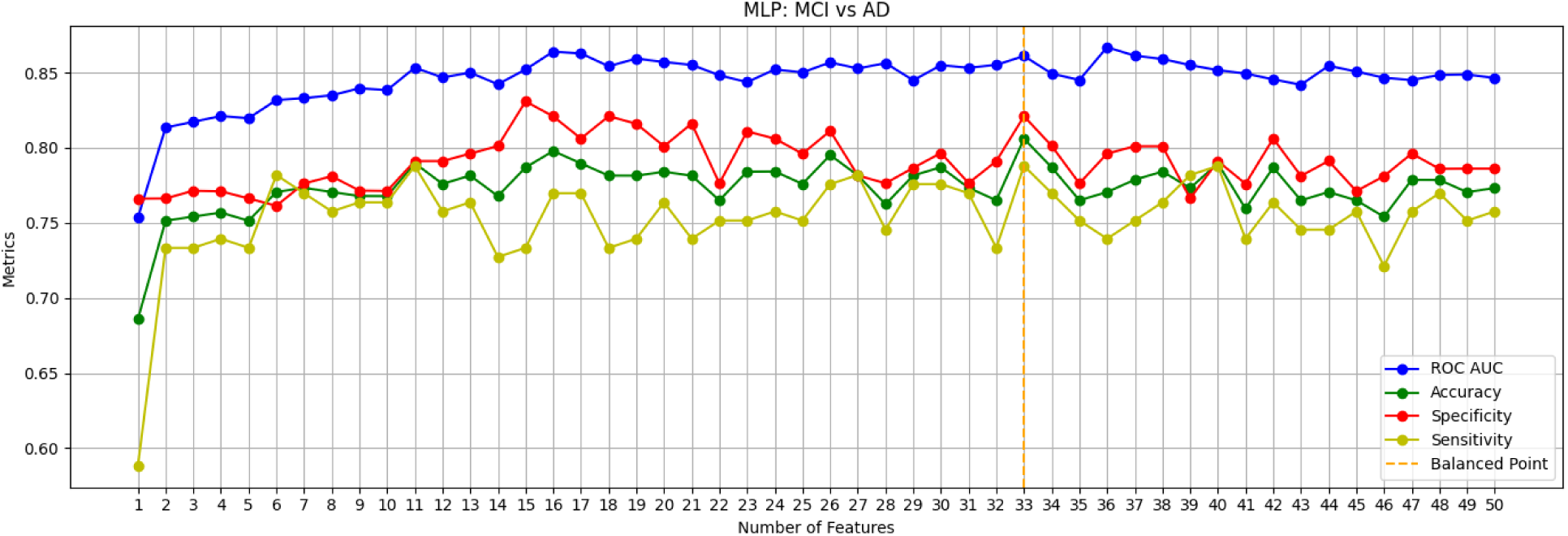
Performance of MLP Classifier with LASSO-Selected Features for MCI vs. AD Discrimination.

To enhance model interpretability and reduce redundancy, Pearson correlation analysis was conducted on the top 33 selected features to identify those with minimal interdependence (Figure 5). This approach prioritized uncorrelated features to ensure that each contributed unique information without overlapping. From this analysis, the feature pairs with the lowest absolute correlation coefficients were selected, reflecting weak or negligible relationships. Their statistical significance was further confirmed using p-values derived from the Spearman rank correlation test, validating the strength of these low correlations and eliminating the likelihood of random variation. The resulting Table 6 presents the selected feature pairs, their correlation coefficients, and corresponding p-values, forming a solid basis for an optimized feature set.

**Fig. 5.**
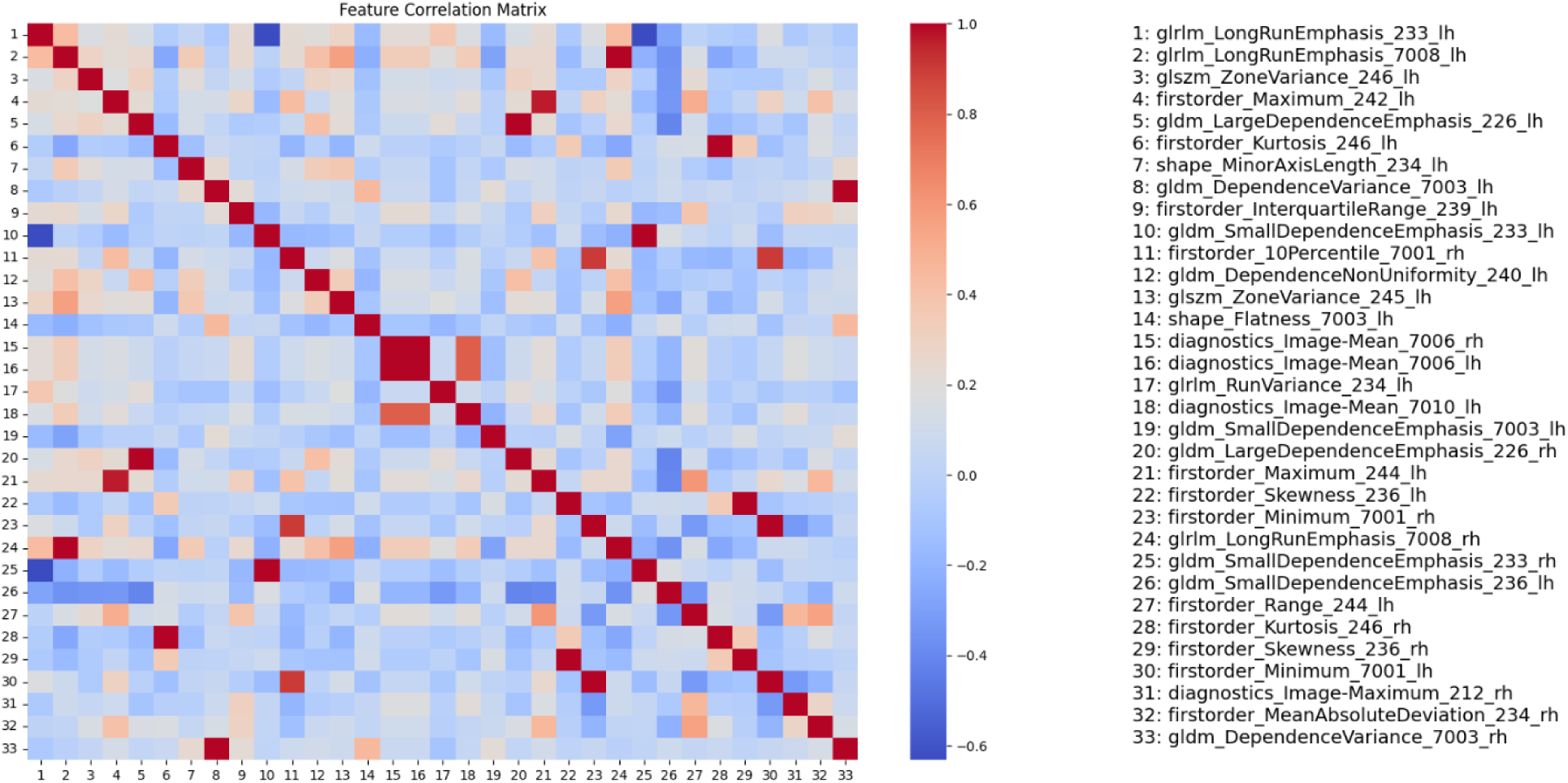
Feature correlation matrix for the group of features with the highest accuracy in discriminating MCI from AD.

**Table 6.** Highly uncorrelated feature pairs for distinguishing MCI vs AD.

| Uncorrelated Features (Feature Name, FS ID, left/right hand) |  | Correlation | P-Value |
| --- | --- | --- | --- |
| original_glrlm_RunVariance_234_lh | original_firstorder_Skewness_236_lh | 0.001 | 5.81E-01 |
| original_gldm_SmallDependenceEmphasis_233_lh | original_glrlm_LongRunEmphasis_7008_lh | 0.002 | 6.72E-01 |
| original_firstorder_Range_244_lh | original_glrlm_LongRunEmphasis_233_lh | 0.002 | 6.55E-01 |
| original_gldm_DependenceNonUniformity_240_lh | original_firstorder_Minimum_7001_lh | 0.003 | 7.80E-01 |
| original_firstorder_MeanAbsoluteDeviation_234_rh | original_glrlm_LongRunEmphasis_233_lh | 0.004 | 3.99E-01 |
| original_gldm_SmallDependenceEmphasis_7003_lh | original_firstorder_Maximum_242_lh | 0.005 | 8.00E-01 |
| original_glrlm_LongRunEmphasis_7008_lh | original_gldm_DependenceVariance_7003_lh | 0.006 | 7.80E-01 |
| original_glrlm_LongRunEmphasis_7008_rh | original_gldm_DependenceVariance_7003_lh | 0.007 | 7.80E-01 |
| original_firstorder_Skewness_236_rh | original_shape_MinorAxisLength_234_lh | 0.008 | 8.05E-01 |
| original_glrlm_RunVariance_234_lh | original_gldm_SmallDependenceEmphasis_7003_lh | 0.009 | 2.09E-01 |

### 3.4 CN vs. MCI

The comparative analysis of CN vs. MCI classification was conducted to identify the most predictive feature subset using the MLP classifier. Similar to the CN vs. AD evaluation, a rank-based approach was applied, systematically evaluating all subsets of the top features identified through the LASSO method. Features were added incrementally based on their ranking by co-efficiency values. For each subset, metrics such as ROC AUC, accuracy, specificity, and sensitivity were computed. The objective was to identify the feature subset that achieved the highest average of ROC AUC and accuracy. Figure 6 illustrates the performance of the MLP classifier for various feature subsets. The optimal subset for CN vs. MCI classification comprised 35 features, achieving an ROC AUC of 0.803, accuracy of 0.731, specificity of 0.735, and sensitivity of 0.726. This configuration was selected as the optimal point based on the average ROC AUC and accuracy, highlighting the effectiveness of the subset in distinguishing between CN and MCI subjects.

**Fig. 6.**
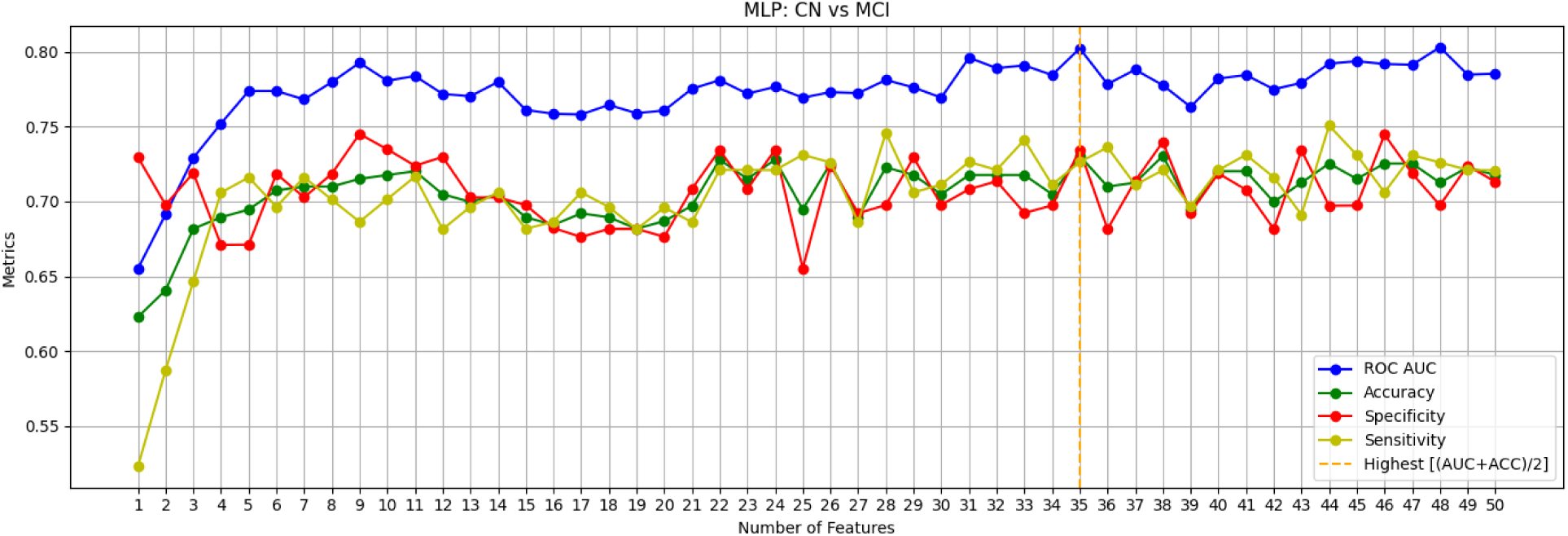
Performance of MLP Classifier with LASSO-Selected Features for CN vs. MCI Discrimination.

To further refine the model and achieve a more interpretable feature set, Pearson correlation analysis was applied to the top selected features (Figure 7). This step aimed to reduce redundancy and ensure the selected features provided unique contributions to the model. By focusing on uncorrelated features, we minimized overlap in the information represented by the selected variables.

**Fig. 7.**
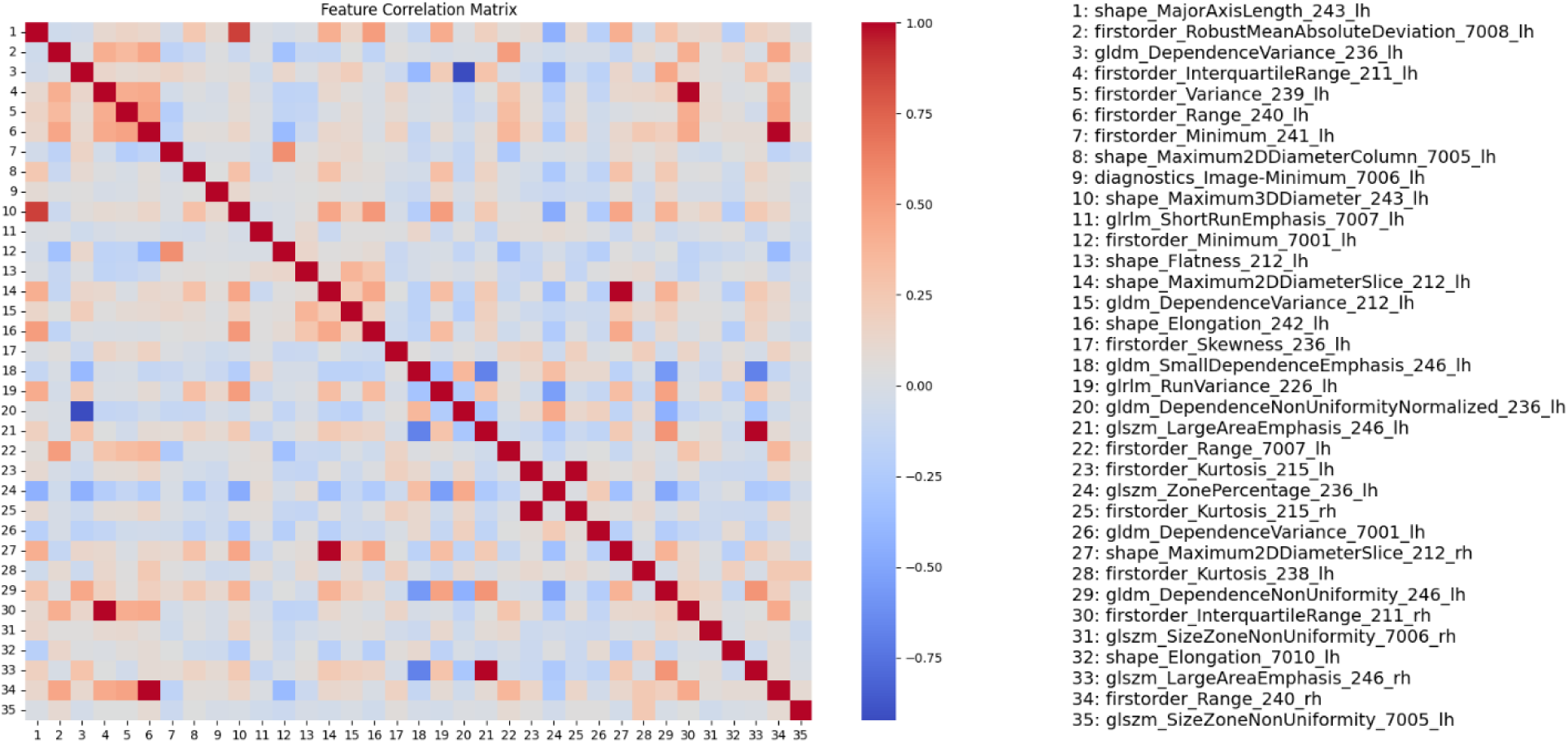
Feature correlation matrix for the group of features with the highest accuracy in discriminating CN from MCI.

To identify the most diverse and representative subset, feature pairs with the lowest absolute correlation coefficients were selected based on Pearson correlation analysis. The statistical significance of these correlations was validated using the Spearman rank correlation test, ensuring the robustness of the feature selection process. The resulting Table 7 summarizes the selected uncorrelated feature pairs, along with their correlation coefficients and corresponding p-values.

**Table 7.** Highly uncorrelated feature pairs for distinguishing CN vs MCI.

| Uncorrelated Features (Feature Name, FS ID, left/right hand) |  | Correlation | P-Value |
| --- | --- | --- | --- |
| original_firstorder_Skewness_236_lh | original_gldm_DependenceVariance_7001_lh | 0 | 0.706 |
| original_firstorder_Range_7007_lh | original_shape_Maximum2DDiameterColumn_7005_lh | 0.001 | 0.7609 |
| original_firstorder_InterquartileRange_211_rh | original_shape_Elongation_242_lh | 0.002 | 0.7156 |
| original_firstorder_InterquartileRange_211_lh | original_shape_Elongation_242_lh | 0.002 | 0.7156 |
| original_firstorder_Kurtosis_215_rh | original_firstorder_Range_7007_lh | 0.002 | 0.5728 |
| original_gldm_DependenceNonUniformityNormalized_236_lh | original_firstorder_RobustMeanAbsoluteDeviation_7008_lh | 0.002 | 0.7097 |
| original_firstorder_Minimum_241_lh | original_firstorder_Minimum_7001_lh | 0.002 | 0.5246 |
| diagnostics_Image-original_Minimum_7006_lh | original_firstorder_Variance_239_lh | 0.002 | 0.8809 |
| original_firstorder_InterquartileRange_211_rh | original_glrlm_RunVariance_226_lh | 0.003 | 0.7854 |
| original_glrlm_RunVariance_226_lh | original_firstorder_InterquartileRange_211_lh | 0.003 | 0.7854 |
| original_glszm_LargeAreaEmphasis_246_lh | original_glszm_SizeZoneNonUniformity_7005_lh | 0.003 | 0.3579 |
| original_firstorder_Variance_239_lh | original_shape_Maximum2DDiameterSlice_212_lh | 0.004 | 0.5479 |

### 3.5 Analysis of Key Features and Regions

Our in-depth analysis of uncorrelated features for both diagnosis groups identified two key features as the common subset for both groups CN vs. AD and MCI vs. AD: Long Run Emphasis (GLRLM) in the left accessory basal nucleus and Small Dependence Emphasis (GLDM) in the left presubiculum-head. These features were found to be the most effective in differentiating between MCI, CN, and AD individuals among the highly uncorrelated feature pairs.

These features achieved remarkable performance in both classification tasks using the MLP classifier. For CN vs. AD, the MLP classifier yielded an ROC AUC of 0.903, accuracy of 0.822, sensitivity of 0.863, and specificity of 0.776. In the MCI vs. AD classification, the same classifier achieved an ROC AUC of 0.824, accuracy of 0.749, sensitivity of 0.776, and specificity of 0.715 as presented in Table 8. Furthermore, for CN vs. MCI, Robust Mean Absolute Deviation (First order) and Minimum (First order) were the most effective uncorrelated features, achieving a ROC AUC of 0.677 and an accuracy of 0.636 when combined.

**Table 8.** Performance evaluation of the two most effective features individually and in combination for classifying CN vs. AD, MCI vs. AD, and CN vs. MCI using MLP.

| Uncorrelated Features |  | ROC AUC | Accuracy | Sensitivity | Specificity |
| --- | --- | --- | --- | --- | --- |
| CN vs. AD | original_gldm_SmallDependenceEmphasis_233_lh | 0.782 | 0.723 | 0.618 | 0.815 |
|  | original_glrlm_LongRunEmphasis_7008_lh | 0.885 | 0.797 | 0.739 | 0.847 |
|  | original_gldm_SmallDependenceEmphasis_233_lh<br>& original_glrlm_LongRunEmphasis_7008_lh | 0.903 | 0.822 | 0.776 | 0.863 |
| MCI vs. AD | original_gldm_SmallDependenceEmphasis_233_lh | 0.748 | 0.686 | 0.558 | 0.791 |
|  | original_glrlm_LongRunEmphasis_7008_lh | 0.798 | 0.735 | 0.691 | 0.771 |
|  | original_gldm_SmallDependenceEmphasis_233_lh<br>& original_glrlm_LongRunEmphasis_7008_lh | 0.824 | 0.749 | 0.715 | 0.776 |
| CN vs. MCI | original_firstorder_RobustMeanAbsoluteDeviation_7008_lh | 0.631 | 0.62 | 0.611 | 0.629 |
|  | original_firstorder_Minimum_241_lh | 0.631 | 0.613 | 0.607 | 0.619 |
|  | original_firstorder_RobustMeanAbsoluteDeviation_7008_lh<br>& original_firstorder_Minimum_241_lh | 0.677 | 0.636 | 0.606 | 0.666 |

Table 9 and Figure 8 together show the structural and textural changes of the hippocampal-amygdala complex across CN, MCI, and AD, underlining the importance of specific radiomic features as biomarkers. In the comparison between CN and AD, the feature Small Dependence Emphasis (GLDM) presents a remarkable growth in mean values, growing from 0.0839 in CN to 0.1636 in AD. This corresponds to a highly increased gray-level heterogeneity. On the other hand, Long Run Emphasis (GLRLM) significantly decreases from 0.6429 to 0.3988, reflecting less structural uniformity. Both features present highly significant p-values, which points out their relevance for diagnosis.

**Table 9.**
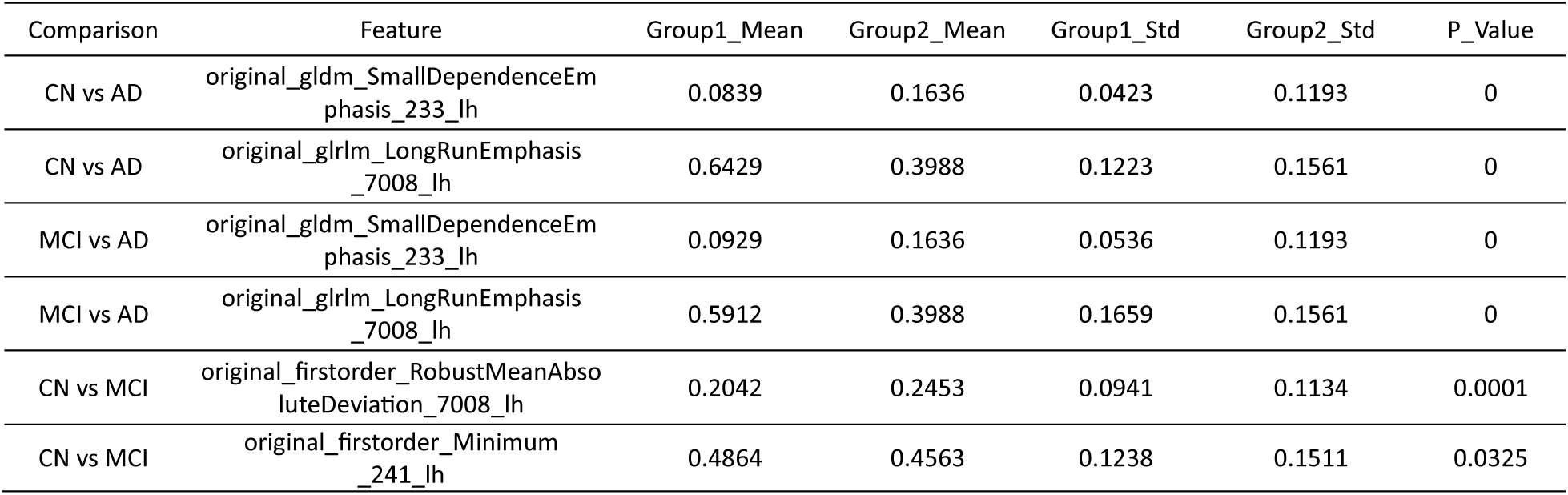
Comparison of Radiomic Features Across CN, MCI, and AD Groups, Highlighting Mean, Standard Deviation, and Statistical Significance.

**Fig. 8.**
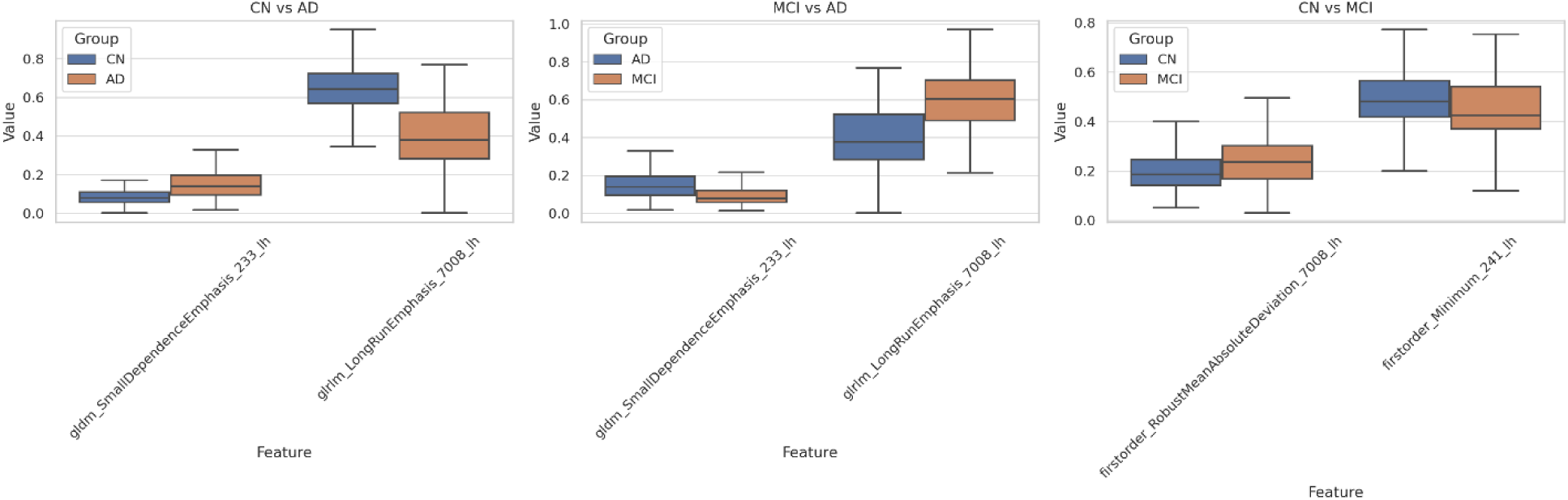
Comparison of Radiomic Features Across Diagnostic Groups: Boxplots illustrate the normalized values of selected radiomic features for three key comparisons: CN vs. AD, MCI vs. AD, and CN vs. MCI.

A similar trend can be seen in MCI vs. AD, where Small Dependence Emphasis (GLDM) increases from 0.0929 in MCI to 0.1636 in AD, and glrlm_LongRunEmphasis decreases from 0.5912 to 0.3988, reinforcing these features as able to tell AD from earlier stages. The changes observed in CN vs MCI are subtler. Robust Mean Absolute Deviation (First order), for example, increases from 0.2042 to 0.2453, indicative of higher variability in the texture in MCI, and the Minimum (First order) decreased from 0.4864 to 0.4563, indicating a loss of signal intensity. This trend is underlined by their respective p-values of 0.0001 and 0.0325.

## 4 Case Study: Longitudinal Analysis of Radiomic Features in AD Progression

### 4.1 Patient Overview

As a case study, we examined the long-term data of a single patient from the ADNI database. Over eight years, the patient was subjected to various scans and clinical assessments, transitioning from CN to AD. We selected four FDG-PET scans that illustrate this progression. Additionally, clinical evaluations were recorded, and scores involving MMSE, ADAS11, ADAS13, and CDRSB scores were presented in Table 10.

**Table 10.**
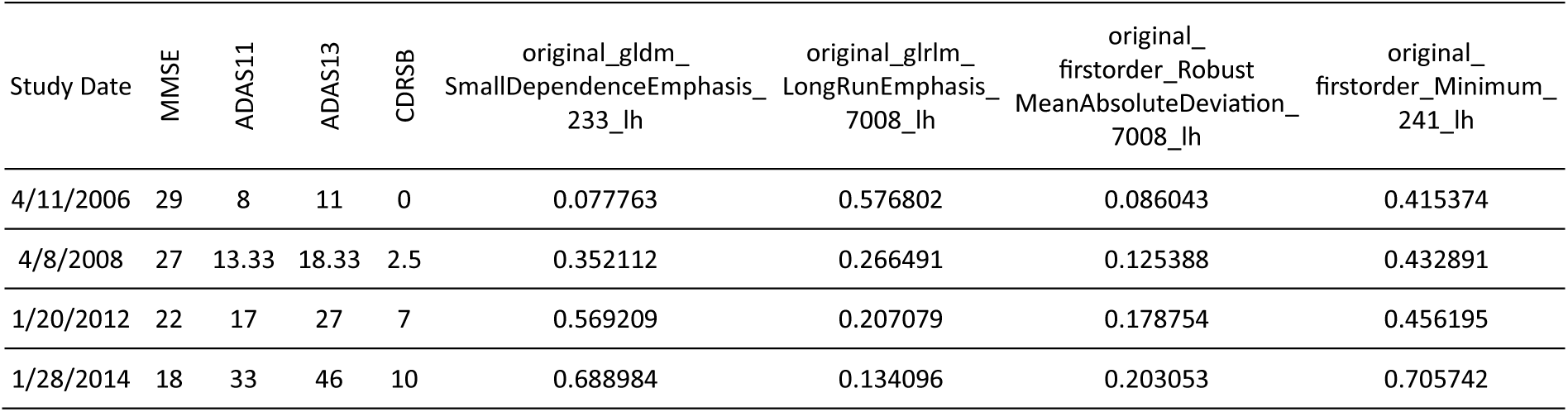
Radiomic features and clinical scores over time.

### 4.2 Methodology

Radiomic features were extracted from the three subregions including the accessory basal nucleus, presubiculum head, and CA4-head using PyRadiomics. Key features such as Small Dependence Emphasis (GLDM), Long Run Emphasis (GLRLM), Robust Mean Absolute Deviation (First order), and Minimum (First order) were computed and analyzed across the four scans.

### 4.3 Results of the case study

The patient exhibited a steady decline in clinical scores (MMSE from 29 to 18 and CDRSB from 0 to 10), correlating with significant changes in radiomic features. The temporal evolution of each radiomic feature is shown in Table 10 and visualized in Figure 9, highlighting trends and abrupt shifts in metabolic patterns. The possible interpretations of observed changes in these computed radiomics features for above mentioned specific three sub-regions of the hippo-amygdala complex are given in the next section.

**Fig. 9.**
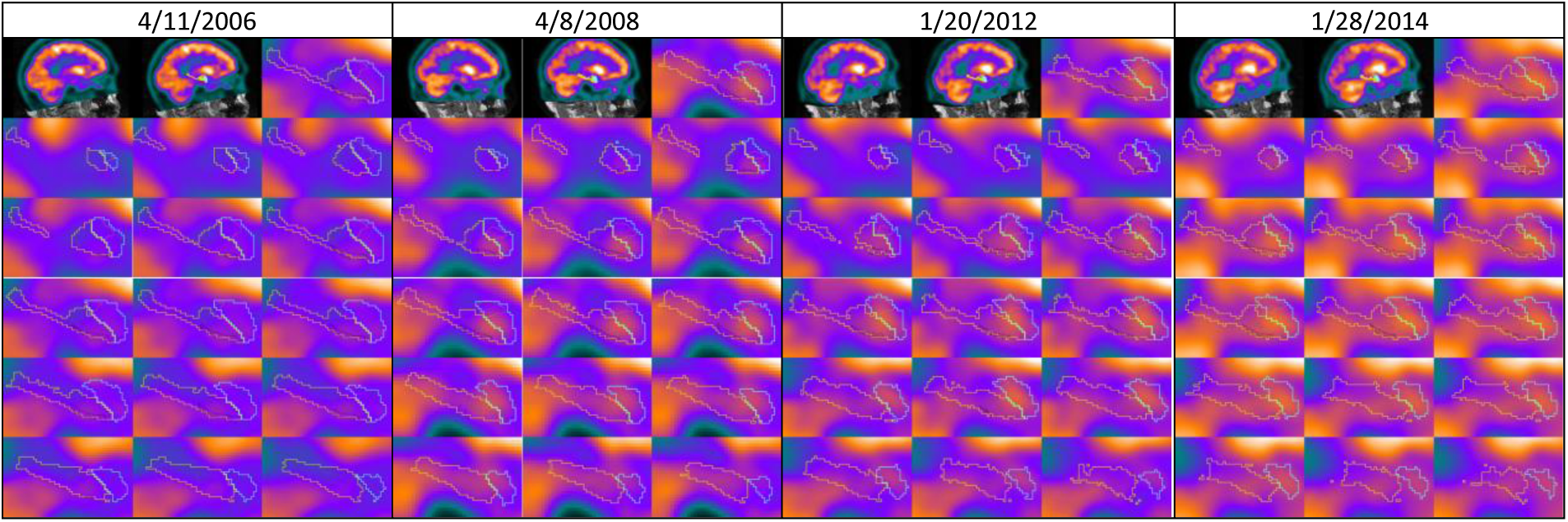
Longitudinal FDG-PET scans with segmented ▪hippocampal and ▪amygdala subregions (▪accessory basal nucleus, ▪Presubiculum, ▪CA4), demonstrating progressive changes in key areas.

## 5 Discussion

This study aimed to identify radiomic features within the hippocampus and amygdala that could differentiate AD from CN and those with MCI. To achieve this, we analyzed baseline FDG-PET scans from 555 participants in the ADNI database. We segmented the hippocampus and amygdala into 38 subfields and 18 subnuclei using probabilistic atlases. We then extracted 120 radiomic features for each segmented subfield and subnucleus using the PyRadiomics tool. By combining various machine learning classifiers and feature selection techniques, we identified the most relevant features for predicting the progression of Alzheimer’s disease. Ultimately, analysis of these features led to a reduction in complexity, resulting in a more interpretable few features. The findings indicate that these radiomic features not only serve as effective diagnostic tools but also offer promise as imaging metabolic biomarkers for early detection and monitoring of AD progression as well as for formulating new hypotheses about the biological mechanism of the disease and defining specific therapeutical targets

The studies summarized in Table 9 explore various imaging and classification techniques targeting Alzheimer’s disease stages, focusing on the hippocampus and amygdala. These studies have all been conducted using MRI imaging. To the best of our knowledge, this is the first study with FDG PET which is known to be highly sensitive due to its molecular nature [39]. It is therefore expected to obtain an earlier diagnosis. It can be seen that the present study presents several novel contributions i) The use of FDG PET imaging ii) The use of a relatively higher number of patients’ iii) The few radiomics features that may help explainability as well as interpretability iv) A relatively high accuracy in discriminating all the stages between each other. The proposed methodology using FDG PET integrates metabolic radiomics features with a Multi-layer Perceptron classifier, achieving superior results (AUC 0.867) in differentiating AD vs. MCI and a competitive AUC (e.g., 0.957 for AD vs. NC) while leveraging a substantial dataset (555 samples). This approach, combining metabolic and regional data, enables early detection and disease monitoring, offering a holistic advantage over solely structural or functional techniques.

**Table 9:** Summary of Imaging Techniques and Methodologies for Classifying Alzheimer’s Disease Stages Based on Hippocampal and Amygdala Features.

| DOI | Methodology | Regions of Interest | Imaging Modality | Dataset Size | Metrics |
| --- | --- | --- | --- | --- | --- |
| [15] | Volumetric analysis using automated segmentation | Hippocampal and Amygdala Subfields | MRI | 92 | AD vs. NC: AUC 0.97<br>MCI vs. NC: AUC 0.79<br>AD vs. MCI: AUC 0.81 |
| [16] | Functional connectivity analysis with support vector machine learning | Hippocampus | fMRI | 119 | AD vs. NC: Acc 0.82<br>MCI vs. NC: Acc 0.81<br>AD vs. MCI: Acc 0.81 |
| [17] | 3D deep convolutional neural network incorporating visual and global shape features | Hippocampus | MRI | 933 | AD vs. CN: AUC 0.978 |
| [18] | Segmentation and volumetric analysis | Hippocampus | MRI | 373 | AD vs. CN: Acc 0.944 |
| Proposed Method | Radiomic feature extraction with Multi-layer Perceptron classifier analysis | Hippocampal and Amygdala Subfields | FDG PET | 555 | AD vs. NC: AUC 0.957<br>MCI vs. NC: AUC 0.782<br>AD vs. MCI: AUC 0.867 |

By concentrating on metabolic data, the study reveals early biochemical changes linked to Alzheimer’s disease, capturing subtle disease progressions that structural imaging might overlook. With a robust sample size of 555 cases, the findings offer strong generalizability and reliability. Given these benefits, our approach provides a framework for both the early identification of Alzheimer’s disease and the differentiation of MCI from AD or normal controls, making it a potential method for clinical and research applications alike.

Our findings identified four significant radiomic features from FDG-PET scans that help distinguish between MCI, AD, and CN states which can potentially capture subtle gray-level variations in brain texture, reflecting underlying Alzheimer’s pathology. Long Run Emphasis (GLRLM), associated with the left accessory basal nucleus, a key area for memory consolidation and emotional regulation, indicates potential functional changes due to neurodegeneration with its increased values.

Small Dependence Emphasis (GLDM), linked to the left presubiculum head involved in spatial navigation and memory, quantifies texture smoothness and detects early metabolic or microstructural changes, possibly preceding visible anatomical alterations like atrophy.

Additionally, Robust Mean Absolute Deviation (First order) and Minimum (First order), are crucial for differentiating CN from MCI. Minimum (First order), tied to the CA4-head region, measures the lowest voxel intensity, signaling subtle shifts in neuronal integrity and metabolic activity that may suggest initial neurodegenerative changes critical for memory encoding.

Meanwhile, Robust Mean Absolute Deviation (First order) assesses the spread of voxel intensities, offering insights into the local heterogeneity of brain tissue. Together, these features provide a comprehensive biomarker for early detection and characterization of Alzheimer’s disease progression.

The case study in Fig 9, Table 10 highlights the potential of radiomic features to detect subtle metabolic shifts that correlate with clinical decline in Alzheimer’s Disease progression:

- An increased Small Dependence Emphasis could suggest that the brain region exhibits a more uniform pattern of glucose metabolism, with less variation in intensity values across neighboring voxels. This could potentially be associated with: More Widespread Neuronal Dysfunction: A more uniform decrease in glucose metabolism across the region, indicating more extensive neuronal damage. Reduced Heterogeneity: A less complex pattern of metabolic activity, potentially reflecting a less dynamic or less active brain region.
- Long Run Emphasis (GLRLM) In the context of FDG PET imaging for Alzheimer’s disease, a decrease in the “Long Run Emphasis” feature from Gray Level Run Length Matrix (GLRLM) analysis might suggest increased heterogeneity in glucose metabolism within the brain regions of interest. Long Run Emphasis: A measure that emphasizes the presence of long runs of pixels with the same intensity. A lower value suggests that there are fewer long runs and more frequent transitions between different intensity levels.

A decrease in Long Run Emphasis (GLRLM) could suggest that the brain region exhibits a more heterogeneous pattern of glucose metabolism, with more frequent transitions between higher and lower glucose uptake areas. This could be due to mixed patterns of neurodegeneration, that is, the presence of areas with more severe neuronal damage alongside areas with less severe damage. Or the presence of inflammatory processes or other metabolic changes that could contribute to increased glucose uptake in certain areas.

- Robust Mean Absolute Deviation (First order) A measure of statistical dispersion that is less sensitive to outliers than the standard deviation. A higher value indicates greater variability in the data. Possible Interpretations:

- Increased Heterogeneity: An increased Robust Mean Absolute Deviation (First order) could suggest that glucose metabolism within the brain region is more variable, with some areas showing higher uptake and others showing lower uptake. This could be due to: i) Mixed Patterns of Neurodegeneration: The presence of areas with more severe neuronal damage alongside areas with less severe damage ii) Inflammation or other Processes: The presence of inflammatory processes or other metabolic changes that could contribute to increased glucose uptake in certain areas.
- Minimum (First order) showed steady increases, contrary to the expected decrease in metabolic activity with disease progression, potentially reflecting inflammation. However, this needs further verification.

Our findings align with existing theories about Alzheimer’s pathology, supporting the view that the hippocampus and amygdala are early targets of neurodegeneration in AD [40], [41]. The distinct textural changes captured by the identified features provide a nuanced characterization of these regions, indicating alterations in gray-level uniformity and structural heterogeneity. By highlighting specific subregions such as the accessory basal nucleus, presubiculum head, and CA4-head, this study demonstrates the potential of radiomics to reveal subtle but clinically significant changes in Alzheimer’s disease progression. These changes correlate with known neurodegenerative processes, including atrophy and disrupted neuronal connectivity [42]. Furthermore, the study goes beyond traditional volumetric measures, offering a more detailed understanding of these brain regions and their involvement in Alzheimer’s pathology.

The findings in Table 8 emphasize, that the accessory basal nucleus may be an important subregion among all subregions of the hippo-amygdala complex for all states of AD. These results highlight the potential importance of the accessory basal nucleus, a subnucleus of the amygdala, in the pathology of Alzheimer’s disease. Its pronounced ability to differentiate diagnostic categories suggests it may play a critical role in early disease mechanisms. Further exploration of the neurobiological factors behind this observation is warranted, as it could deepen our understanding of Alzheimer’s progression, particularly regarding functional changes in the amygdala. The findings in this study confirm the important knowledge about metabolic changes in the hippo-amygdala complex in Alzheimer’s disease

Our evaluation framework meticulously examined the robustness and performance of the model through a structured analysis involving nine classifiers combined with four feature selection techniques, resulting in 36 distinct configurations. Each configuration underwent rigorous evaluation based on two key metrics—accuracy (ACC) and area under the ROC curve (AUC)—to comprehensively assess predictive performance. This detailed approach highlights the reliability and adaptability of our methodology, culminating in the selection of the MLP classifier coupled with LASSO for feature selection. To ensure unbiased results, stratified k-fold cross-validation (k=5) was implemented. The selected models exhibited strong performance in distinguishing between CN and AD (ROC AUC: 0.957, Accuracy: 0.907), MCI and AD (ROC AUC: 0.867, Accuracy: 0.806), and CN and MCI (ROC AUC: 0.782, Accuracy: 0.753).

There are several limitations of our study. First, the cross-sectional design of the FDG-PET data limits the assessment of the longitudinal trends component intrinsic to furthering the understanding of the disease progression [43]. Second, the generalizability of the findings may be limited by the peculiar characteristics of the ADNI cohort. Increasing the number and diversity of populations and databases used for data extraction will enhance the robustness and applicability of the identified biomarkers [44]. Furthermore, even though feature reduction enhanced the interpretability of the biomarkers, it may have unintentionally excluded features of potential relevance, particularly those reflecting subtle but critical changes in other brain regions that are highly correlated with the selected features. This limitation underscores the need for careful consideration of biologically meaningful correlations during feature selection to avoid discarding valuable information. Thus, this result needs further investigation in independent data with a view to their validation and further integration of radiomic biomarkers with clinical and genetic data for refining predictive models. To facilitate the clinical translation of these findings, it is crucial to address challenges such as standardization of image acquisition protocols, feature extraction methods, and analysis pipelines. Additionally, larger, multicenter studies are needed to validate the generalizability of these findings and to develop robust clinical decision support systems [45].

In conclusion, this study highlights the promise of FDG PET radiomic biomarkers of specific subregions for the early diagnosis and management of Alzheimer’s disease. By focusing on the hippocampus and amygdala, we provide novel possible explanations for the disease’s metabolic underpinnings. Using only a few numbers of features for staging can help the interpretability of this scheme. Future efforts should concentrate on the effects of tailored targeted therapies on the discovered biomarkers and refine further their biological interpretation.

## Acknowledgments

We would like to acknowledge the support from the Boğaziçi University Research Fund (BAP) under project code 19774, which played a critical role in facilitating this research.

Data collection and sharing for this project was funded by the Alzheimer’s Disease Neuroimaging Initiative (ADNI) (National Institutes of Health Grant U01 AG024904) and DOD ADNI (Department of Defense award number W81XWH-12-2-0012). ADNI is funded by the National Institute on Aging, the National Institute of Biomedical Imaging and Bioengineering, and through generous contributions from the following: AbbVie, Alzheimer’s Association; Alzheimer’s Drug Discovery Foundation; Araclon Biotech; BioClinica, Inc.; Biogen; Bristol-Myers Squibb Company; CereSpir, Inc.; Cogstate; Eisai Inc.; Elan Pharmaceuticals, Inc.; Eli Lilly and Company; EuroImmun; F. Hoffmann-La Roche Ltd and its affiliated company Genentech, Inc.; Fujirebio; GE Healthcare; IXICO Ltd.; Janssen Alzheimer Immunotherapy Research & Development, LLC.; Johnson & Johnson Pharmaceutical Research & Development LLC.; Lumosity; Lundbeck; Merck & Co., Inc.; Meso Scale Diagnostics, LLC.; NeuroRx Research; Neurotrack Technologies; Novartis Pharmaceuticals Corporation; Pfizer Inc.; Piramal Imaging; Servier; Takeda Pharmaceutical Company; and Transition Therapeutics. The Canadian Institutes of Health Research is providing funds to support ADNI clinical sites in Canada. Private sector contributions are facilitated by the Foundation for the National Institutes of Health (www.fnih.org). The grantee organization is the Northern California Institute for Research and Education, and the study is coordinated by the Alzheimer’s Therapeutic Research Institute at the University of Southern California. ADNI data are disseminated by the Laboratory for Neuro Imaging at the University of Southern California.

